# Electrophysiological evidence of synergistic auxin transport by interacting Arabidopsis ABCB4 and PIN2 proteins

**DOI:** 10.1101/2021.01.25.428116

**Authors:** Stephen D. Deslauriers, Edgar P. Spalding

## Abstract

To understand why both ATP-binding cassette B (ABCB) and PIN-FORMED (PIN) proteins are required for polar auxin transport through tissues, even though only the latter is polarly localized, we biophysically studied their transport characteristics separately and together by whole-cell patch clamping. ABCB4 and PIN2 from *Arabidopsis thaliana* expressed in human embryonic kidney cells displayed electrogenic activity when CsCl-based electrolytes were used. Current-voltage (I-V) analysis of the activities and modeling the effects of adding the auxin anion (IAA^−^) as a potential substrate with the Goldman-Hodgkin-Katz equation, demonstrated that ABCB4 and PIN2 were 9-fold and 10-fold more selective for IAA^−^ than Cl^−^, respectively. Thus, these proteins directly transport IAA^−^, which was not unequivocally established by previous auxin retention assays. Co-expression of ABCB4 and PIN2 produced an especially significant result. Co-expression synergistically doubled the selectivity for IAA^−^. An area of two-fold higher selectivity for IAA^−^ that this result indicates will occur in cells with asymmetric PIN2 and symmetric ABCB4 matches what early models found to be necessary to create observed levels of polar auxin transport through tissues. Thus, the requirement for two different proteins appears to be explained by a synergistic effect on selectivity. More substrate details and important pharmacological results are reported.

## Introduction

A special mechanism for directing auxin to its sites of action was recognized even before the chemical identity of this important plant hormone was known (Went and Thimann, 1937). So-called polar auxin transport was originally proposed to result from channels at the downstream faces of each cell releasing auxin anions down a large thermodynamic gradient (Rubery and Sheldrake, 1974; Raven, 1975; Goldsmith, 1977). Discovery of PIN-FORMED (PIN) proteins that are required for polar auxin transport and asymmetrically localized at the plasma membrane in accordance with the theory was a genuine breakthrough (Okada et al., 1991; Chen et al., 1998; Gälweiler et al., 1998; Křeček et al., 2009; Petrášek et al., 2006). However, it became apparent that a PIN-only explanation was insufficient when members of the B subfamily of ATP-Binding Cassette (ABCB) transporters were also shown to be essential for polar auxin transport yet not asymmetrically localized in the cells performing the transport (Sidler et al., 1998; Noh et al., 2001; Terasaka et al., 2005; Cho et al., 2007; Wu et al., 2007; Wu et al., 2010). Why asymmetrically-localized PIN proteins and symmetrically-localized ABCB proteins are both essential for polar auxin transport is an open question (Spalding, 2013).

The most widespread view of PIN and ABCB function, based largely on radioactively-labelled auxin retention assays, is that both types of proteins are auxin transporters (Geisler et al., 2005; Petrášek et al., 2006; Yang and Murphy, 2009; Kubeš et al., 2012; Kamimoto et al., 2012). Tests of the prevailing view with independent methodologies would be valuable because alternative explanations of the existing data are plausible (Spalding, 2013). The principal method for studying electrogenic transporters, those that produce an electric current when transporting their substrate, is to express the transporter in a non-plant cell and then use patch-clamp electrophysiology to analyze any ionic currents attributable to the expressed transporter (Dryer et al., 1998). This approach demonstrated that the ABCB19 protein from Arabidopsis produced ion channel activity with weak selectivity for anions over cations when the human embryonic kidney (HEK) cell was the heterologous expression system (Cho et al., 2014). The anion channel blocker that Noh et al. (2001) used in the screen for upregulated genes that originally identified *ABCB19*, a chemical called 5-nitro-2-(3-phenylpropylamine)-benzoic acid (NPPB), blocked the ABCB19 channel activity in HEK cells. The same low concentration of NPPB also blocked polar auxin transport very effectively in Arabidopsis roots, and blocked gravitropism, indicating that the anion channel activity in the heterologous system was functionally relevant.

The Arabidopsis *ABCB19* gene was originally called *MDR1* (Noh et al., 2001) because when it was isolated, the only similar sequence in the database was human *MDR1*, also called *P-gp1*, and now known as ABCB1. Human ABCB1 is best known for its role in removing chemotherapeutic molecules from tumor cells (Ambudkar et al., 2003). It has also been reported to possess inorganic ion transport activity, or to modulate a separate ion transport activity (Valverde et al., 1992; Higgins, 1995; Hoffman et al., 1996; Roepe, 2000; Fletcher et al., 2010). The more distantly related ABCC7 protein, called the cystic fibrosis transmembrane regulator (CFTR), is a Cl^−^ channel with a pore that is blocked by NPPB (Wang et al., 2005; Csanády and Töröcsik, 2014). In the case of plant ABCB transporters, the auxin anion (IAA^−^) may be a natural substrate, transported across the membrane in a channel-like fashion. The blocking effects of NPPB on the channel and polar auxin transport (Cho et al., 2014), and the impaired polar auxin transport in *abcb* mutants (Noh et al., 2001; Lewis et al., 2007) support this idea. Still lacking is a demonstration of an ABCB protein transporting IAA^−^ channel-like, i.e. passively in the thermodynamic sense. In the case of PIN proteins, their role as auxin transporters is inferred from the amount of auxin retained in cells engineered to express or lack them (Geisler et al., 2005; Petrášek et al., 2006; Yang and Murphy, 2009; Weller et al., 2017;). No electrophysiological investigations of PIN transport activity have been reported. Presumably, they transport IAA^−^ passively, i.e. thermodynamically downhill and therefore should show electrogenic activity. Some of the experiments reported here directly test if ABCB and PIN proteins electrogenically transport IAA^−^ in the manner of thermodynamically passive efflux channels.

To understand why both ABCB and PIN are both necessary for the polar auxin transport phenomenon in tissues, their interactions must be understood in addition to their separate transport activities. Co-immunoprecipitation and yeast two-hybrid assays indicate PIN and ABCB proteins physically interact. Evidence of functional interaction comes from measurements of radioactive auxin retained in cells expressing both protein types (Yang and Murphy, 2009; Blakeslee et al., 2007; Mravec et al., 2008; Titapiwatanakun et al., 2008; Cho et al., 2012). It has been suggested (Spalding, 2013) that interactions between the two proteins may produce a synergistic function, potentially explaining why polar auxin transport is disrupted in a mutant that cannot place ABCB transporters in the plasma membrane but properly places PIN transporters (Wu et al., 2010; Wang et al., 2013).

The experiments reported here used ABCB4 and PIN2 because these members of the two protein families required for auxin transport are expressed in the same outer cell layers of the Arabidopsis root, both are required for shootward polar auxin transport, and both participate in the root gravitropism response (Abas et al., 2006; Lewis et al., 2007). They are a logical pair of proteins to study separately and together with new methods in order to advance our understanding of the polar auxin transport mechanism.

## Results

### ABCB4 and PIN2 display weakly anion-selective channel activity

To test the hypothesis that ABCB4 and PIN2 transport activities are electrogenic and therefore may be studied biophysically with the patch clamp technique, cDNA encoding one or the other of these proteins was expressed in cultured HEK cells. The bi-cistronic transfection vector also encoded a fluorescent protein marker in order to determine which cells in a field were suitable for patch clamp analysis. The major component of the pipette and bath solutions was 140 mM CsCl, chosen to preclude currents from endogenous sodium and potassium channels. In response to a voltage-step protocol, cells expressing ABCB4 or PIN2 displayed time-independent inward and outward currents that were three or four-fold greater than currents recorded from controls cells transfected with a vector containing only the fluorescent marker (Figure 1A). Current-voltage (I-V) relationships in these symmetric CsCl conditions were approximately linear and reversed (I=0) at −5 or −8 mV (Figure 1B). A rigorous method to measure relative ion selectivity is to change the concentration difference between the pipette and bath solutions and then measure the shift in reversal potential of the I-V curve (Hille, 2001). Reducing the CsCl concentration in the bath solution from 140 mM to 14 mM to create an asymmetric condition shifted the reversal voltage (E_rev_) of control cells by −11 mV. The Goldman-Hodgkin-Katz (GHK) equation (Equation 1) relates E_rev_ to the membrane’s permeability to the principal ions present as explained by Hille (2001).

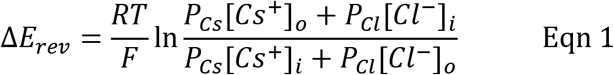

**Figure 1.**
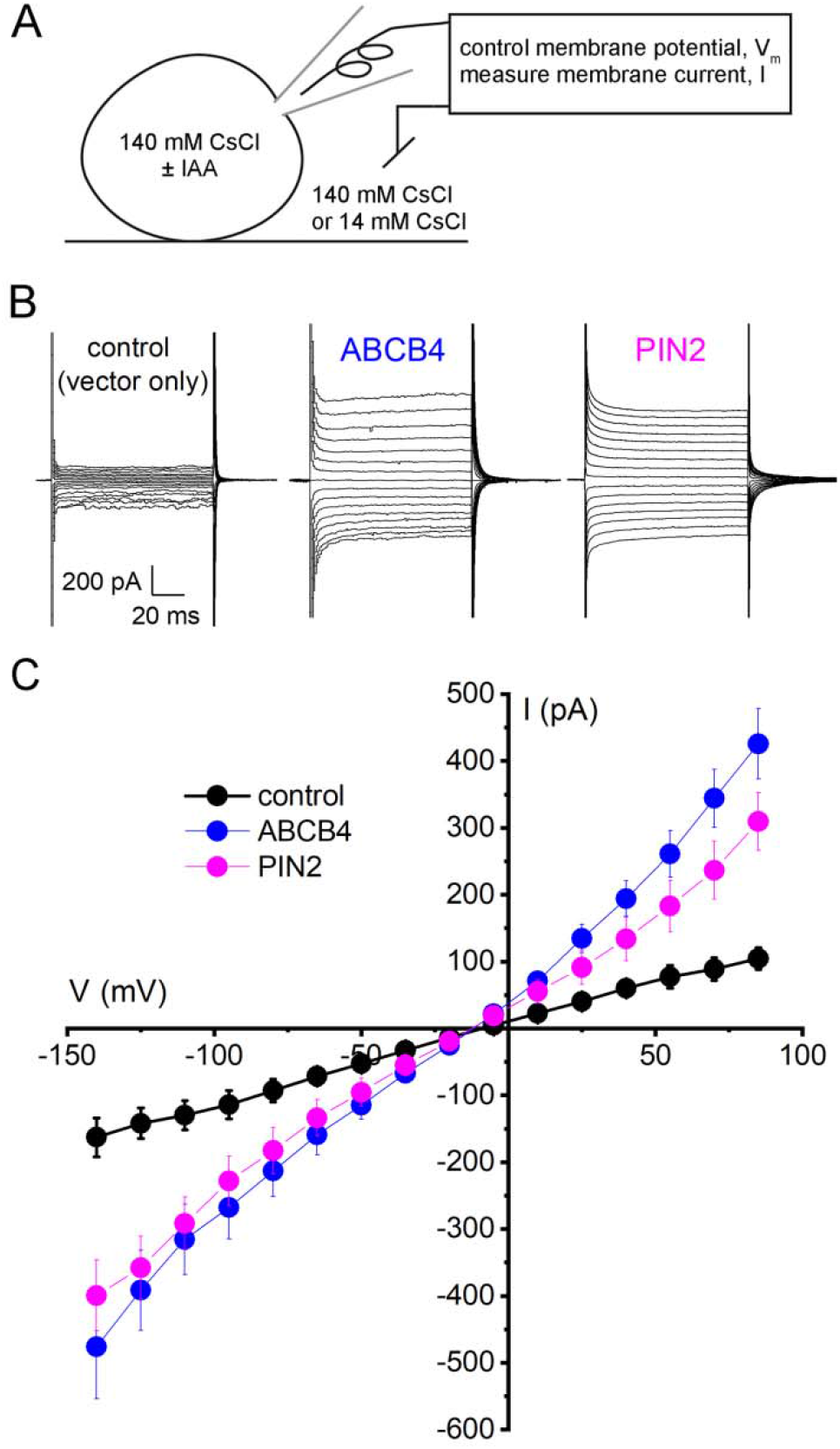
Electrogenic activities of ABCB4 and PIN2 proteins expressed in HEK cells. **A**, Diagram of the whole-cell patch clamp technique employing Cs^+^ and Cl^−^ as principal charge carriers. The amplifier controlled (clamped) the membrane potential (V) while the transmembrane electric currents (I) were measured. The cells were transfected with a vector carrying only EGFP or DsRED cDNA (control), ABCB4 and EGFP cDNA in separate reading frames, or PIN2 and DsRED cDNA in separate reading frames. **B**, Example recordings of transmembrane currents elicited by step-wise changes in V recorded from control cells and cells transfected with ABCB4 or PIN2. **C**, I versus V curves represent mean current ± SE at each membrane potential measured in control cells (n = 5), ABCB4-expressing cells (n = 6), and PIN2-expressing cells (n = 6). The pipette and bath solutions contained 140 mM CsCl.

A ΔE_rev_ of −11 mV indicates the average control plasma membrane was 0.59 times as permeable to Cl^−^ as it was to Cs^+^ (P_Cl_:P_Cs_ = 0.59). The same shift of the bath solution from 140 mM to 14 mM CsCl shifted E_rev_ of ABCB4-expressing cells significantly in the opposite direction, to 11 mV (Figure 2A), corresponding to a P_Cl_:P_Cs_ = 1.7. The plasma membrane of PIN2-expressing cells displayed a ΔE_rev_ of 23 mV, corresponding to a P_Cl_:P_Cs_ = 3.1 (Figure 2B). Thus, expressing ABCB4 or PIN2 shifted the HEK cell membrane from less to more permeable to Cl^−^ relative to Cs^+^, consistent with a role in auxin transport because essentially all indole-3-acetic acid is in the anionic form (IAA^−^) at the pH of cytoplasm (Raven, 1975; Spalding, 2013). When ABCB4 and PIN2 were co-expressed, the P_Cl_:P_Cs_ of the membrane was 2.5 (Figure 2C). To separate the the activities of the plant proteins from background permeabilities, the average endogenous HEK cell (control) I-V curve was subtracted from the average experimental I-V curves, the E_rev_ for each difference curve was determined, and the P_Cl_:P_Cs_ for the expressed transporter was calculated. Figure 2D shows the difference I-V curves and the calculated P_Cl_:P_Cs_ for ABCB4 (2.1), PIN2 (3.6), and ABCB4+PIN2 (5.1).

**Figure 2.**
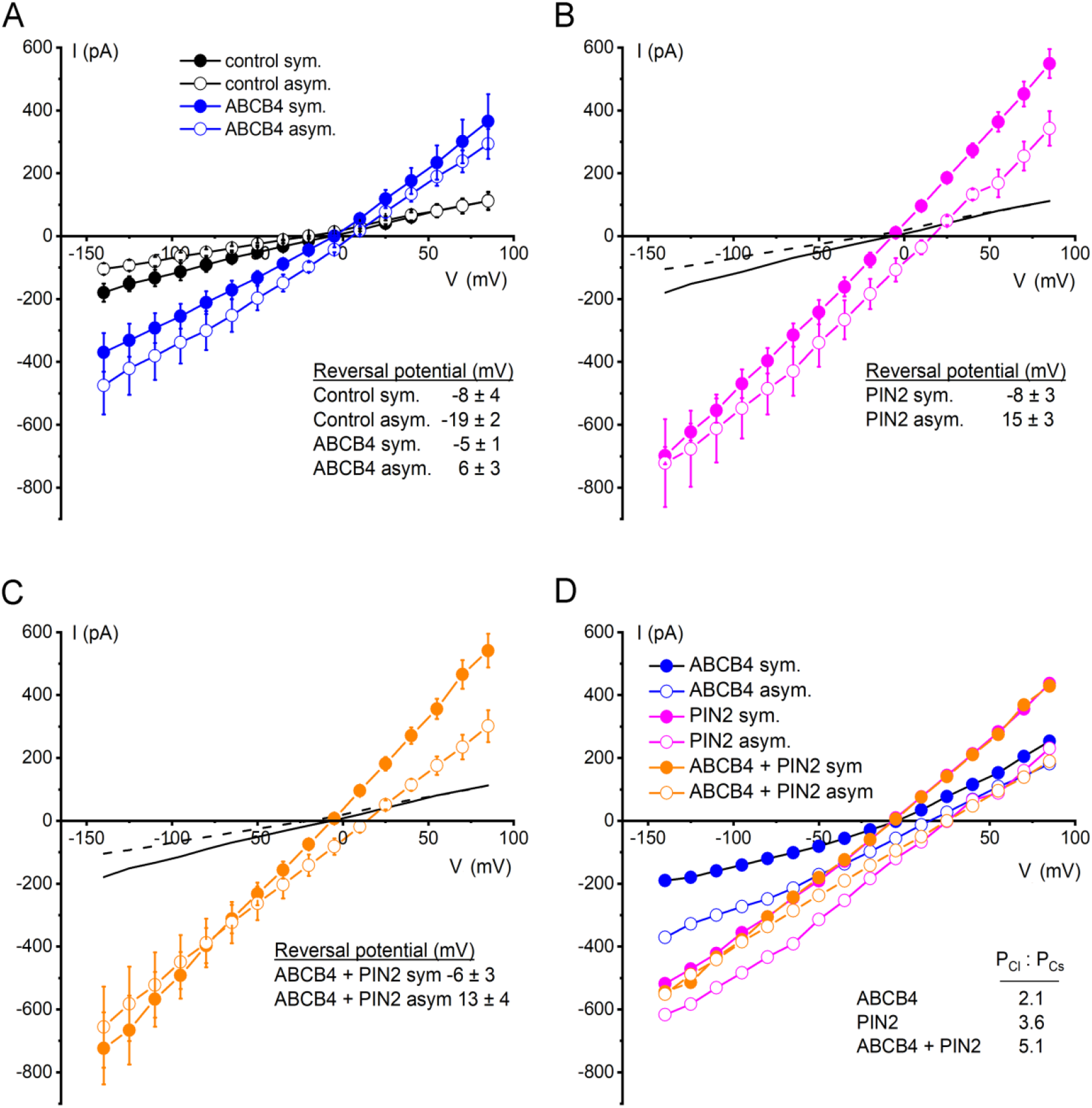
Anion preferences ABCB4, PIN2, and co-expressed ABCB4 and PIN2 transport activities demonstrated by current-voltage analysis. **A**, ABCB4 and control cell I-V relationships recorded with 140 mM CsCl in the bath and pipette (symmetrical) and after switching the bath to 14 mM CsCl (asymmetrical). The average membrane potentials at which I=0 (reversal potential) for each condition are shown. Plotted are the mean currents ± SE at each voltage obtained from 4 separate cells for each condition. Positive changes in reversal potential indicate preference for Cl^−^ over Cs^+^. **B**, same as A but for cells expressing PIN2. The control cell curves are re-plotted from A. **C**, same as B but for cells co-expressing ABCB4 and PIN2. **D**, ABCB4-, PIN2-, and ABCB4+PIN2-dependent I-V curves were generated from the data in A-C by subtracting average control currents from each. The Cl^−^ to Cs^+^ permeability ratios (P_Cl_:P_Cs_) derived from the reversal potentials of these curves are included in the plot.

### ABCB4 and PIN2 conduct IAA^−^

To test whether ABCB4 or PIN2 can also conduct IAA^−^, auxin in the pipette solution was increased from 0.1 μM to 1 mM IAA in a solution containing only 50 mM CsCl to reduce background or competing currents (Figure 3A,B). If ABCB4 or PIN2 proteins conduct IAA^−^, increasing its concentration in the pipet should shift E_rev_ to a more positive voltage. Figure 3C,D shows that significant positive shifts were observed in cells expressing ABCB4, PIN2, and ABCB4+PIN2, but not in control cells. These biophysical measurements establish that ABCB4 and PIN2 transport IAA^−^ across the membrane. Furthermore, the physiological relevance of the different magnitudes of the IAA-dependent shifts in E_rev_ become apparent when the results are analyzed with a GHK-based model of the experiment (Eqn 2).

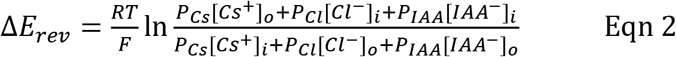

**Figure 3.**
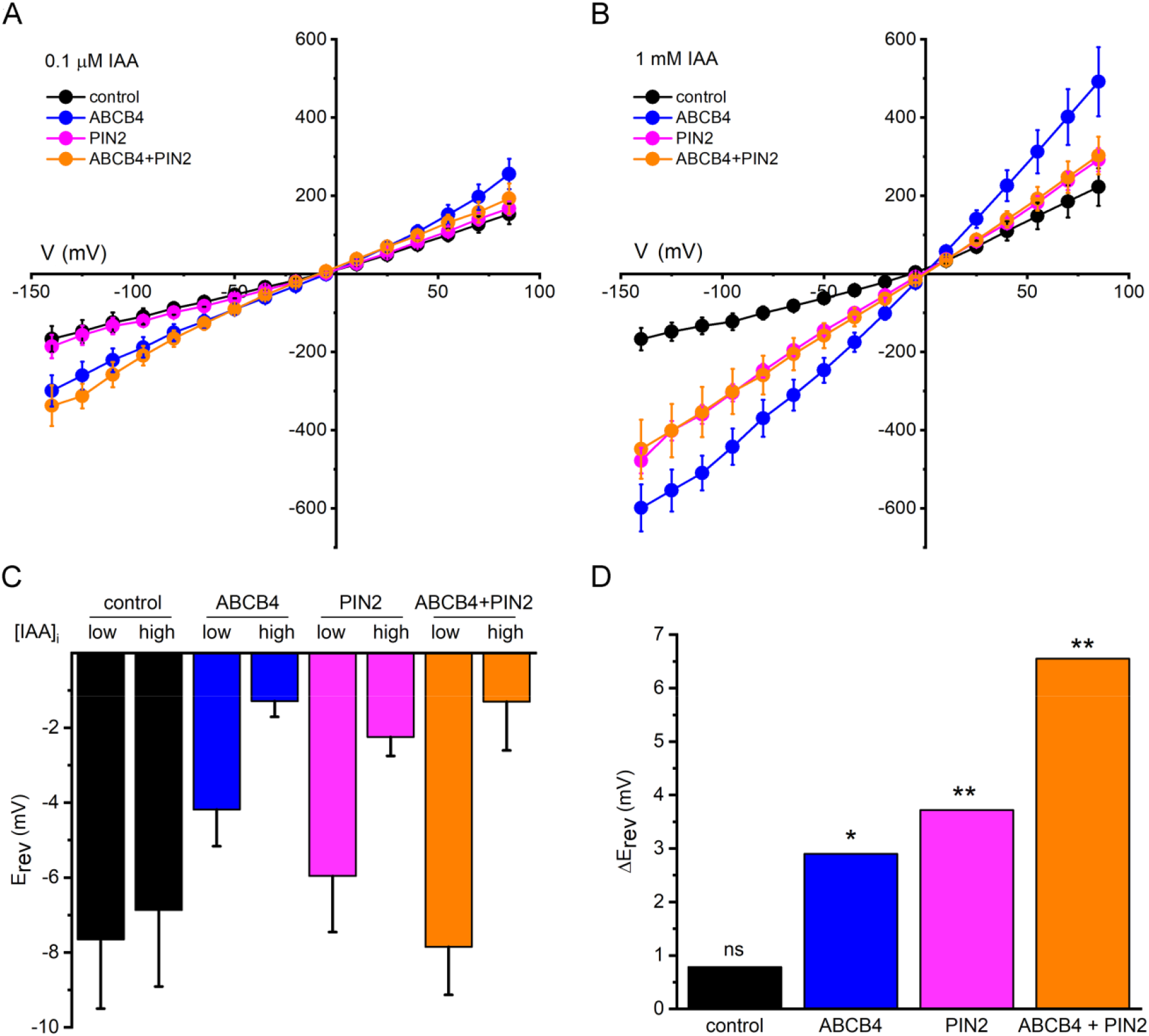
IAA^−^ permeability demonstrated by current-voltage analysis. **A**, I-V curves obtained in symmetrical 50 mM CsCl conditions with 0.1 μM IAA in the pipette, i.e. on the cytoplasmic side. **B**, I-V curves obtained as in A but with 1 mM IAA in the pipette. The number of independent cells measured per condition was between 4 and 14. **C**, Reversal potentials (E_rev_) of I-V curves obtained with either 0.1 μM IAA (low) or 1 mM IAA (high) in the pipette. **D**, Differences in E_rev_ (ΔE_rev_) due to a change in the IAA gradient. T-tests were performed. ns = no statistical significance, * = p<0.05, ** = p<0.01.

The IAA-dependent shifts in E_rev_ can be used to determine P_IAA_ relative to P_Cl_ for each of the transporters and their combination (Figure 4). The values used to parameterize the GHK model (Eqn 2) were derived from the data in Fig. 2D. P_Cl_:P_Cs_ was 2.1 for ABCB4, 3.6 for PIN2, and 5.1 for ABCB4+PIN2. The resulting curves show that the measured auxin-dependent E_rev_ shifts (Figure 3D), i.e. 2.9 mV for ABCB4, 3.8 mV for PIN2, and 6.6 mV for ABCB4+PIN2, reflect a P_IAA_:P_Cl_ of 9 for ABCB4, 10 for PIN2, and 18 for ABCB4+PIN2. Thus, a membrane containing ABCB4 and PIN2 are manifold more selective for IAA^−^ than for Cl^−^, and the activity formed by the combination of these proteins is approximately 2-fold more selective for IAA^−^ over Cl^−^ than either protein alone. The results in Figures 3 and 4 constitute biophysical evidence of ABCB4 and PIN2 proteins directly transporting IAA^−^, and that selectivity of the permeation pathway(s) for IAA^−^ approximately doubled when both proteins were present in the same membrane. The results show that co-expression did not increase the activity. In fact, the amount of ionic current flowing at each of the imposed membrane potentials was reduced compared to ABCB4 expressed alone (Figure 3B). Rather, the contribution of IAA^−^ to the transported current is what roughly doubled when the two proteins were co-expressed.

**Figure 4.**
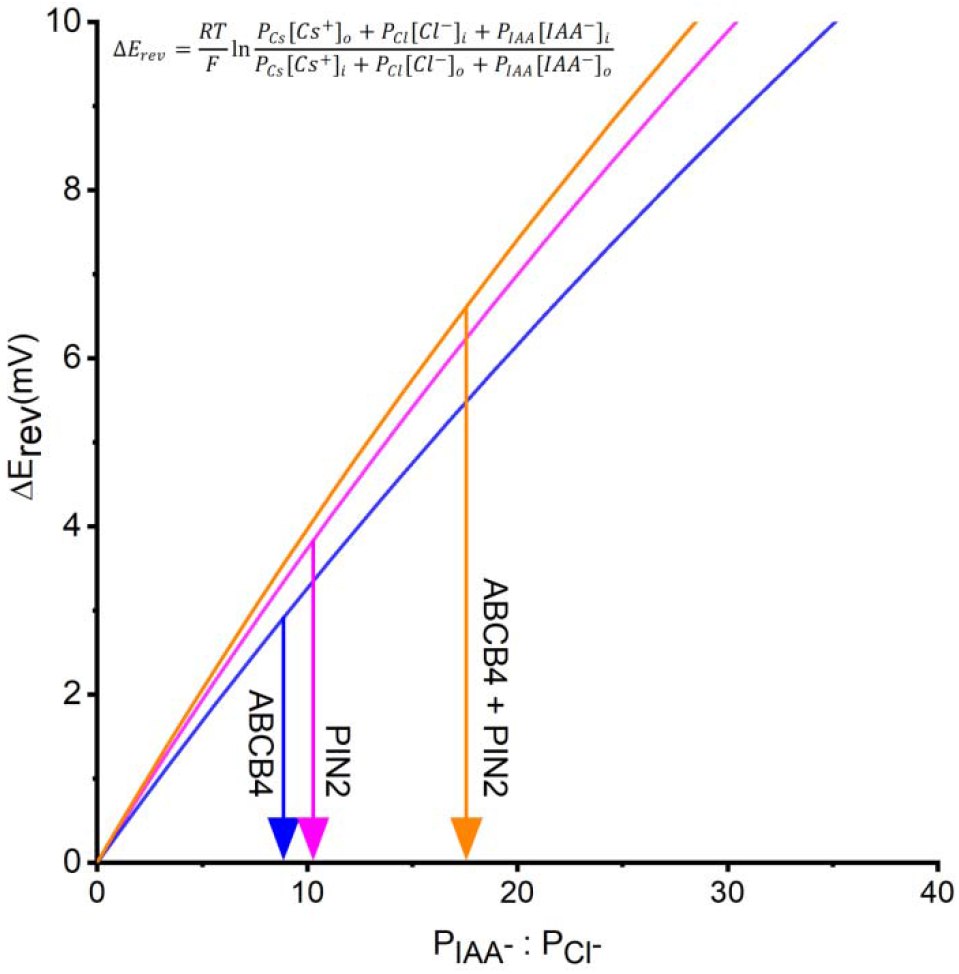
ABCB4 and PIN2 are highly selectivity for IAA^−^ over Cl^−^, and the selectivity approximately doubles when the two are co-expressed. The Goldman-Hodgkin-Katz model of membrane transport was parameterized with values of (P_Cl_) relative to Cs^+^ permeability (P_Cs_) calculated from the results in Figure 2D to determine the permeability of IAA^−^ relative to Cl^−^ (P_IAA_ : P_Cl_) based on the ΔE_rev_ values presented in Figure 3D.

While testing whether IAA^−^ transport could be detected with electrophysiology methods, a stimulatory effect of the hormone on transporter activity was observed, particularly for ABCB4 (compare Figure 3A and Figure 3B). Additional experiments were performed to investigate the potential for auxin regulation of the auxin transport activity. Figure 5A shows that increasing IAA concentration in the pipette from 0.1 μM to 1 μM increased the current in ABCB4-expressing cells. Increasing it further to 1 mM further increased the activity (data from Figure 3B). PIN2 activity was also stimulated by auxin but not within the presumed physiologically relevant range of 0.1-1 μM (Figure 5B). Benzoic acid (BA) is a weak acid often used as a non-transported control compound in polar auxin transport assays. It did not stimulate ABCB4 or PIN2 channel activity (Figure 6A,B). Auxin (indole-3-acetic acid) is structurally related to the amino acid tryptophan (Trp), which did not stimulate activity like auxin (Fig. 6A,B). These results demonstrate that IAA but not any weak acid or related indole compound activates ABCB4 and PIN2 transport.

**Figure 5.**
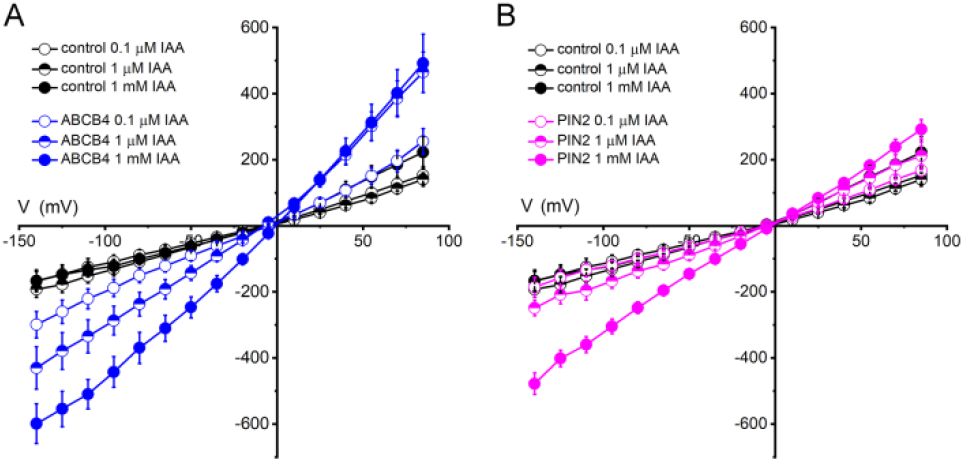
Effects of auxin on transport activity of ABCB4 and PIN2. **A**, I-V relationships for ABCB4 obtained with different concentrations of IAA^−^ in the pipette. **B**, I-V curves for PIN2 obtained with different concentrations of IAA^−^ in the pipette. The pipette and bath solutions contained 50 mM CsCl. The same control data are plotted in A and B, the 0.1 μM IAA and 1 mM IAA data are replotted from Figure 3. The 1 μM IAA data are mean currents ± SE obtained from n= 6 control cells, n=4 ABCB4 cells, and n=5 PIN2 cells.

**Figure 6.**
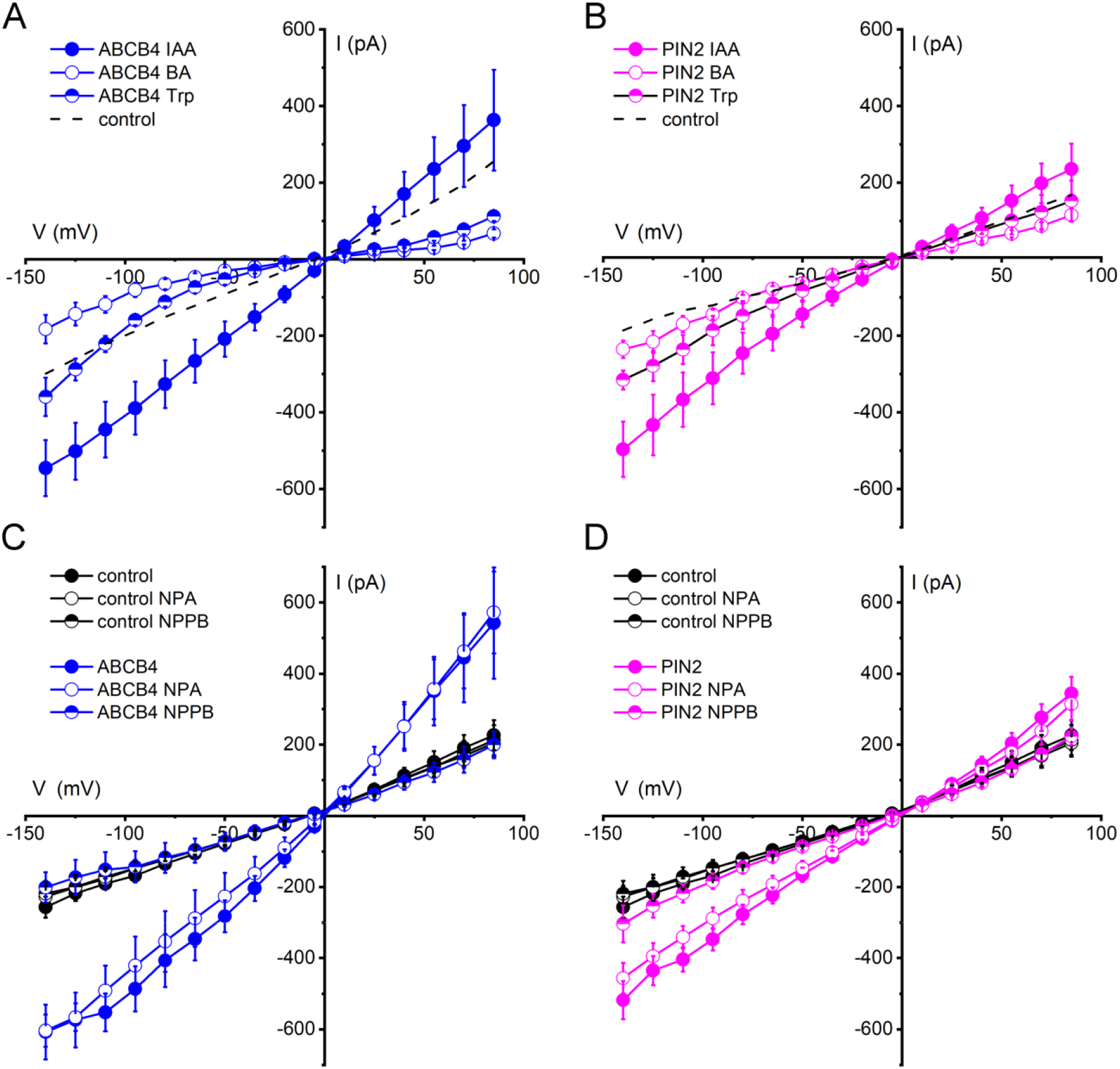
Activation specificity and pharmacology of ABCB4 and PIN2 activity. **A**, The indole-based amino acid Trp nor the aromatic benzoic acid (BA) activates ABCB4 similarly to IAA. The dashed line shows the 0.1 μM IAA ABCB4 baseline copied from Figure 5A. **B**, Same as A but for PIN2. The dashed line shows the 0.1 μM IAA PIN2 baseline copied from Figure 5B. **C**, ABCB4 activity in auxin-stimulated conditions is blocked by 20 μM NPPB but not 10 μM NPA applied by switching the bath solution. Neither treatment significantly affected the control cell currents. **D**, Same as C but for PIN2. Plotted is mean current ± SE at each voltage for n = 4 or 6 independent cells for each treatment. The pipette and bath contained 50 mM CsCl. Untreated control cell data are replotted from Figure 5A.

The experiments designed to test whether Trp or BA could stimulate ABCB4 or PIN2 transport activity also tested whether these compounds sometimes used as control treatments in auxin research were transport substrates. They were not. E_rev_ did not significantly shift when 0.1 μM IAA was replaced with 10^4^-fold higher concentrations (1 mM) of Trp or BA (Supplemental Figure 1). The proteins transported IAA^−^ but not Trp or BA.

### NPPB but not NPA inhibits ABCB4 and PIN2 activity

NPPB is used to block anion channels including those encoded by mammalian ABC transporters (Wang et al., 2005; Csanády and Töröcsik, 2014), and it reversibly blocks Cl^−^-permeable channels in the Arabidopsis plasma membrane (Cho and Spalding, 1996). It displays a half-inhibition concentration of approximately 5 μM (Noh and Spalding, 1998). The Arabidopsis *ABCB19* gene was originally isolated by screening for NPPB-induced genes (Noh et al., 2001). ABCB19 was shown to possess NPPB-inhibited activity when expressed in HEK cells and studied with the patch-clamp methods used in the present study (Cho et al., 2014). NPPB also blocks polar auxin transport in roots as effectively as null *abcb19* mutations (Cho et al., 2014). Consistent with these results, 20 μM NPPB completely blocked ABCB4 channel activity (Figure 6C). The chemical *N*-1-naphthylphthalamic acid (NPA) is widely used in the low micromolar range to inhibit polar auxin transport and has been reported to bind to ABCB19 (Kim et al., 2010) but it did not inhibit ABCB19 activity assayed by patch clamping in HEK cells (Cho et al., 2014). Figure 6C shows that 10 μM NPA did not inhibit ABCB4 activity either. The same pharmacological profile was observed for PIN2. The anion-channel blocker NPPB but not NPA inhibited PIN2-mediated currents (Figure 6C,D). The lack of an inhibitory effect of NPA on PIN2-mediated currents carrying IAA^−^ is consistent with a recent report (Abas et al., 2021) that NPA binds to PIN proteins at a dimerization interface, suggested to be “distinct from IAA substrate-binding sites” and therefore probably not part of the transport pathway within the protein. HEK cells possess an anion channel that could potentially confound the heterologous expression approach used in the present study. However, the HEK cell must be swelled in hypotonic conditions to observe this endogenous activity and 100 μM NPPB is required to suppress it (Hélix et al., 2003). Therefore, this endogenous volume-regulated anion channel (VRAC) is an unlikely contributor to the present results, which we ascribe to the expression of ABCB4 or PIN2.

The Supplemental Dataset 1 contains the 191 individual I-V data sets, each an independent trial obtained from a separate HEK cell, used to generate the results in Figures 1–6.

### Biophotonic assay of ABCB4-PIN2 interaction

The increase in IAA^−^ selectivity detected when ABCB4 and PIN2 were co-expressed may be the result of a physical interaction. In order to investigate a potential interaction between ABCB4 and PIN2 in a live-cell system, fluorescent ABCB4-CFP and PIN2-YFP fusion proteins were co-expressed in HEK cells and tobacco (*Nicotiana benthamiana*) leaf epidermal cells. Figure 7 shows that the efficiency of Förster resonance energy transfer (FRET) from the CFP to the YFP, which requires molecular-scale proximity of the two fluorophores, was greater than that obtained with free CFP and YFP. The efficiency of FRET between ABCB4 and PIN2 shown in Figure 7 is low compared to FRET between subunits comprising amino-acid gated Ca^2+^ channels known as GLRs recorded in the same experimental system (Vincill et al., 2013). Nonetheless, the FRET efficiency was significantly above background in both plant and animal cell membranes, and therefore provides some support to the electrophysiological evidence of interaction (Figures 2–4). FRET between ABCB4 and PIN2 was not enhanced by treating the cells with IAA. Treatment with NPPB may have weakened the interaction between the two proteins as the FRET signal in NPPB-treated leaves was not different from the free CFP/YFP control.

**Figure 7.**
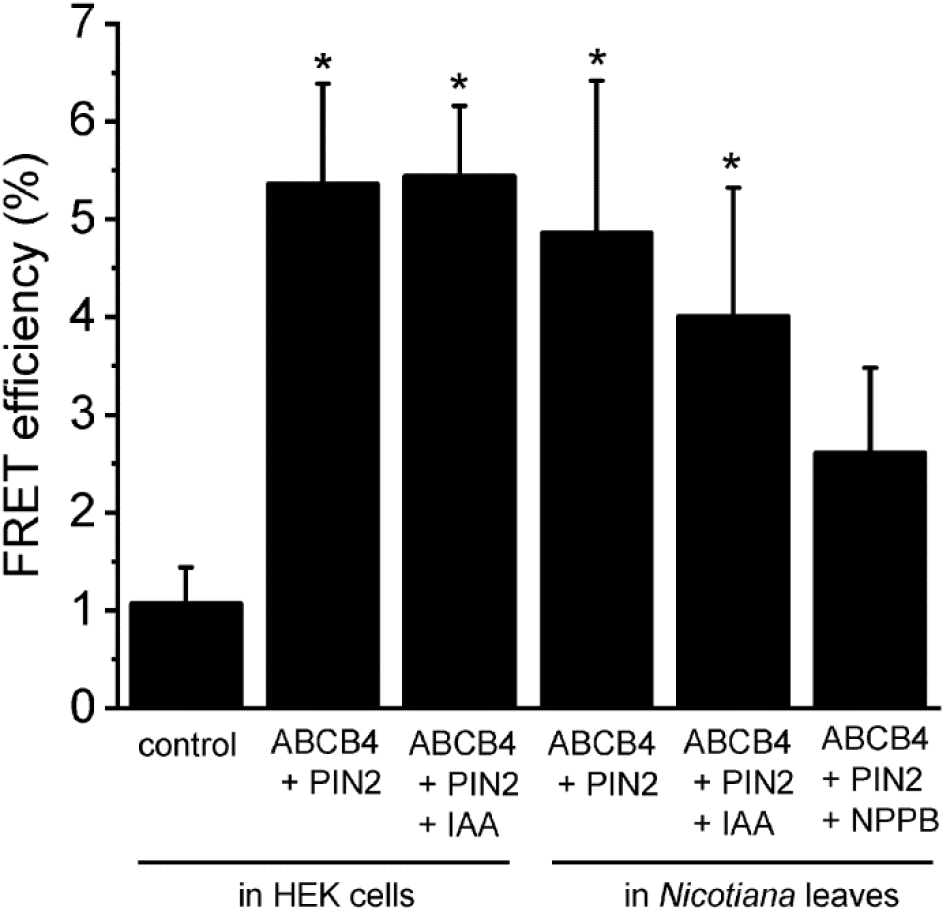
FRET assay of interaction between ABCB4 and PIN2 co-expressed in HEK cells or *Nicotiana benthamiana* leaf epidermal cells. Mean FRET efficiencies ± SE were quantified after photobleaching the acceptor. Control (n = 7), ABCB4-CFP:PIN2-YFP in HEK cells (n = 14), ABCB4-CFP:PIN2-YFP in HEK cells with 1 mM IAA (n = 24), ABCB4-CFP:PIN2-YFP in *N. benthamiana* (n = 8), B4-CFP:PIN2-YFP in *N. benthamiana* with 1 mM IAA (n = 6) and ABCB4-CFP:PIN2-YFP in *N. benthamiana* with 20 μM NPPB (n = 4). Asterisks indicate values that are different to a statistically significant degree from the control (p = 0.05) as determined by T-tests. The control represents the amount of FRET between free CFP and YFP.

Supplemental Figure 2 shows an image of a HEK cell expressing ABCB4-CFP and PIN2-YFP obtained during the course of a FRET assay. The figure indicates the plasma membrane region in which FRET was quantified and the effects of photobleaching the YFP acceptor. Supplemental Dataset 2 shows the pre-bleach and post-bleach fluorescence intensities from each of the fluorophores for every trial that generated data shown in Figure 7. The data show that ABCB4-CFP and PIN2-YFP signal levels were similar, indicating similar concentrations of the two proteins at the membrane. The data also show that in most cases, fluorescence intensity from the donor (CFP) increased following photobleaching of the acceptor (YFP), which is an indicator of genuine FRET.

## Discussion

Mathematical modeling indicated that the difference in IAA^−^ permeability between opposite cell ends required to produce the measured rate and directional bias of auxin transport through oat coleoptiles should be approximately two-fold (Goldsmith et al., 1981; Mitchison, 1981). Figures 3,4 and 7 present evidence that co-expressed ABCB4 and PIN2 interact, to some extent physically, to create a conductance that selects IAA^−^ approximately two-fold better than does either protein separately. Ionic currents conducted by the combination of ABCB4 and PIN2 would be enriched two-fold for IAA^−^ relative to currents conducted by either single protein. This enrichment would be localized to regions of PIN2 expression, leading to polarized efflux of auxin from each cell and thus directionally biased auxin flow through the tissue.

A mechanism for polarized auxin efflux that depends on ABCB and PIN proteins combining to produce an enhanced function would explain why the *twisted dwarf 1* mutation, which prevents ABCB proteins from reaching the plasma membrane (Wu et al., 2010), disrupts polar auxin transport even though PIN protein localization is normal (Bouchard et al., 2006; Wang et al., 2013). The mechanism proposed here also explains why individual *abcb* mutations impair polar auxin transport as severely as *pin* mutations (Gälweiler et al., 1998; Noh et al., 2001; Lewis et al., 2007). Certainly, PINs direct auxin flow in plants (Wiśniewska et al., 2006). The present results explain why ABCB proteins are also needed.

Many important details remain to be discovered or clarified. A better understanding of the nature of the interaction between ABCB and PIN proteins may indicate how the pair selects better for IAA^−^. The FRET efficiency measurements (Figure 7) indicate that the two proteins were close enough often enough to allow a resonance process to transfer excitation energy between them. More and different types of experiments are needed to determine if a complex of ABCB and PIN proteins is an accurate model for the efflux transporter that produces the polar auxin transport phenomenon. A physical interaction between ABCB19/PGP19 and PIN1 strong enough to enable coimmunoprecipitation of the pair was reported (Blakeslee et al., 2007). Mravec et al. (2008) based on different evidence concluded that ABCBs and PINs “interact intermolecularly at the PIN-containing polar domain.” Thus, multiple lines of evidence support a functional and apparently physical interaction between ABCB4 and PIN2.

Until heterologously expressed and studied by patch clamp electrophysiology, it was not known that plant ABCB proteins could conduct ions across the membrane. The first demonstration of such activity was achieved with Arabidopsis ABCB19. Because this activity was in some respects novel, it was important to test for explanations based on artifacts. For example, could ABCB19 in the membrane of the HEK cell have increased the activity of a channel native to the HEK cell? Arguing against this possibility is that expressing ABCB19 containing a missense mutation known to create a null phenotype in Arabidopsis did produce any ion transport activity in the HEK cell (Cho et al., 2014). Co-expressing ABCB19 and the TWD1 chaperone, rather than leading to greater activity as hoped, produced none (Cho et al., 2014). These results indicate that the plant protein did not in some nonspecific way cause the HEK cell to produce a new activity much larger than background. The present study adds more reasons to trust that the activities measured are not artifacts. For example, ABCB4 activity was specifically enhanced by auxin, which would not be expected of an endogenous HEK cell transporter. The transporters are more selective for IAA^−^ than Cl^−^, Trp, or BA (Figures 3,4; Supplemental Figure 1) which would not be expected of a mammalian transporter. These are reasons to trust that the expression system produces a valid representation of the plant protein’s natural function. Therefore, it is necessarily the case that ABCB4 and PIN2 transport numerously more inorganic ions such as Cl^−^ per unit time across the plant cell plasma membrane than IAA^−^ because the concentration of auxin (~μM) is at least one thousand-fold lower than the cytoplasmic concentrations of Cl^−^ and other potentially permeant substrates (~mM). Thus, IAA^−^ is among the ions the auxin efflux machinery allows to move thermodynamically ‘downhill’ across the plasma membrane.

Another plant membrane protein that transports an organic anion also transports Cl^−^. It is the ALMT1 malate transporter, which is activated by Al^3+^ and negatively regulated by gamma-aminobutyric acid (Piñeros et al., 2008; Ramesh et al., 2015). As more plant proteins expected to transport organic compounds are studied electrophysiologically in heterologous systems, more may be found to also transport inorganic ions. The inevitably mixed-ion current conducted by an auxin-efflux activity comprised of ABCB4 and PIN2 may be physiologically important. It may provide cells with a physiological mechanism for sensing IAA^−^ flux, which is a parameter in some models of auxin transport-mediated development (Kramer, 2008; Stoma et al., 2008; Prusinkiewicz et al., 2009; Shinohara et al., 2013) despite no generally recognized notion of how a cell could measure it. A mixed ion current would depolarize the plasma membrane, i.e. shift the membrane potential to more positive values. If interaction between ABCB and PIN increases selectivity for IAA^−^, the depolarizing effect of nonselective currents may be reduced. The downstream (PIN-containing) end of an auxin-transporting cell may have a more negative membrane potential than the upstream end. This cellular electrical polarity will be proportional to the magnitude of polar auxin flux. Spatial differences in membrane potential can drive current loops, long associated with morphogenesis (Jaffe and Nuccitelli, 1977; Hotary and Robinson, 1990; Léonetti et al., 2004; Adams and Levin, 2013). We suggest that electrical consequences of an auxin efflux activity comprised of ABCB and PIN proteins could form the basis of an auxin flux sensor at the cellular, tissue, and organ levels. Manipulation of PIN localization followed by investigations of microscopic and macroscopic electrical gradients could test this idea.

Auxin stimulation of ABCB4 activity, especially if it occurs in other members of the ABCB family, would also contribute to the positive reinforcement of auxin transport by auxin transport itself. Self-strengthening auxin transport is a key feature of canalization, the phenomenon thought to create auxin distribution patterns that guide many aspects of development (Stoma et al., 2008; Sauer et al., 2006; Bennett et al., 2014). The present findings of auxin-stimulated ABCB4 transport activity and increased selectivity due to ABCB4 and PIN2 interactions would augment auxin promotion of PIN expression that is believed to result in canalization (Kramer, 2008; Bayer et al., 2009; Smith and Bayer, 2009; van Berkel et al., 2013).

The results reported here establish a new category of evidence indicating that ABCB4 and PIN2 directly transport IAA^−^ across a cell membrane. The results are based on voltage-clamp measurements of charge movement and membrane transport theory. Because the patch clamp method controls the fundamental parameters governing transport while it directly measures charge movement, it could be the most appropriate method for investigating structure-function relationships of auxin transporters. Other ABCB proteins to be studied with this method include ABCB1, ABCB6, ABCB19, and ABCB20 because they have each been shown to play a role in auxin transport through tissues (Noh et al., 2001; Yang et al., 2018). One structure-function question to address is whether or not the proposed IAA binding sites in ABCB4 (Yang et al., 2009) are responsible for auxin activation of its transport function (Figure 5A). Another is the role of predicted NPA binding sites (Kim et al., 2010) in channel function. The measurement platform could also be used to investigate the regulatory effects of co-expressed kinases (van Berkel et al., 2013; Zourelidou et al., 2014; Weller et al., 2017). Especially important will be biophysical investigations of how apparently new properties such as enhanced selectivity result from interaction of ABCB4 and PIN2, which may be the basis of the synergistic auxin transport Blakeslee et al. (2007) reported. The phenomenon may be analogous to the interaction between mammalian ABCC8 and the Kir6 (K_ATP_) potassium channel which results in regulation not evident in either individual protein (Bryan and Aguilar-Bryan, 1999; Burke et al., 2008). All of the above would benefit from improvements to the platform such as increasing expression of the plant proteins in HEK cells and finding ways to achieve greater than 1 mM IAA in the pipette. The former would increase the activity to be measured and the latter would create larger shifts in E_rev_ and therefore increase the precision with which IAA^−^ selectivity could be measured. The resulting improved understanding of how auxin transport proteins create directionally biased, self-reinforcing auxin flow would lead to more accurate mechanistic models of how auxin influences plant growth and development.

## Materials and Methods

### HEK cell culture and transfection

HEK293T cells from American Type Culture Collection were cultured in Dulbecco’s modified Eagle’s medium-GlutaMAX (Invitrogen) with 10% fetal bovine serum, 100 IU mL^−1^ penicillin, and 100 μg mL^−1^ streptomycin in an incubator set at 37 °C with 95% air and 5% CO_2_. Prior to transfection, trypsin-treated HEK293T cells (5 × 10^5^ cells per well) were plated into six-well tissue culture plates containing collagen-coated glass cover slips 12 to 24 h before being transfected with 1 μg of the indicated plasmid DNA using FuGENE 6 transfection reagent (Promega) following the manufacturer’s protocol. In the case of cotransfections, a plasmid ratio of 1:1 (0.5 μg + 0.5 μg) was used. All experiments were performed 36 to 48 h after transfection and imaged live at room temperature.

### DNA cloning

For HEK293T cell expression and electrophysiology experiments, full length *ABCB4* and *PIN2* cDNA was amplified from total RNA by RT-PCR as described in Vincill et al. (2012) using the following primers: 5’-ATCTGTCGACATGGCTTCAGAGA GCGGCTTA-3’ and 5’-AATTCCCGGGTCAAGAAGCCGCGGTT-3’. PCR products were then digested and inserted into the SalI and XmaI sites of the pIRES-Enhanced Green Fluorescent Protein (EGFP) bicistronic vector used by Vincill et al. (2012) such that a single mRNA would separately code ABCB4 and EGFP. PIN2 was amplified from cDNA template using the following primers: 5’-TATTTGTCGACATGATCACCGGCAAA GAC- 3’ and 5’-ATCACCCGGGTTAAAGCCCCAAAAGAAC-3’. PCR products were digested and inserted into the SalI and XmaI sites of the pIRES2-DsRed bicistronic vector.

For FRET analysis in HEK293T cells, *ABCB4* cDNA was amplified using the following primers to include the CACC Gateway modification on the forward primer: 5’-<u>CACC</u>ATGGCTTCAGAGAGCGGC-3’ and 5’-AGAAGCCGCGGTTAGATGAAGC-3’. PIN2 cDNA was amplified using the following primers: 5’-<u>CACC</u>ATGATCACCGGCAAAGAC ATGTAC-3’ and 5’-AAGCCCCAAAAGAACGTAGTACAGTAC-3’. The resulting PCR fragments were cloned into the pENTR-D entry vector (Invitrogen). The pENTR-D vectors containing the respective full-length *ABCB4* or *PIN2* cDNAs were shuttled into the pDS_EF1-XB-CFP (ABCB4) and pDS_EF1-XB-YFP (PIN2) mammalian expression vectors from American Type Culture Collection using the Gateway recombination reaction (Invitrogen) to generate C-Terminally tagged fusion proteins.

For FRET experiments in *Nicotiana benthamiana*, the pENTR-D vectors described above containing *ABCB4* and *PIN2* cDNA were shuttled into the Gateway destination vectors pEARLEYGATE 101(CFP) and pEARLEYGATE 102(YFP) respectively, which fuse the indicated fluorescent tag to the C-terminus of the translated gene product to generate Pro35S:PIN2-YFP and Pro35S:ABCB4-CFP constructs. The tobacco leaf infiltration method was used to transiently express the constructs. All constructs were confirmed by DNA sequencing.

### Electrophysiology

For whole-cell recording, a coverslip with cells was placed in a recording chamber mounted on the fixed stage of an upright fluorescence microscope (Olympus BX51WI) mounted on an antivibration table equipped with a micromanipulator that controlled the head stage of the patch-clamp amplifier (Axopatch 200A; Molecular Devices;www.moleculardevices.com). A 40x dipping objective lens was used to view the cells in bright-field or fluorescence mode in the chamber, which was being continuously perfused with a bath solution containing 140 mM CsCl, 2 mM CaCl_2_, 2 mM MgCl_2_, 5 mM KCl, and 10 mM HEPES, adjusted to pH 6 with CsOH. The pipette was filled with 140 mM CsCl, 1 mM CaCl_2_, 2 mM MgCl_2_, 5 mM EGTA, 10 mM D-Glc, 10 mM HEPES, and 3 mM Mg-ATP, adjusted to pH 7.2 with CsOH. For experiments using 50 mM CsCl, 190 mM sorbitol was added to maintain osmolarity. Bath solutions containing only 14 mM CsCl solutions were supplemented with 252 mM sorbitol. Cells displaying strong EGFP or DsRed fluorescence were selected for whole-cell patch-clamp analysis using micropipettes pulled from borosilicate glass. Micropipette resistance was between 5 and 8 megaohms when filled. After achieving a gigaohm seal, the patch was ruptured to obtain the whole-cell configuration. After the baseline current stabilized, a voltage clamp protocol was administered by pCLAMP 10.2 software (Molecular Devices). The measured membrane currents were low-pass filtered at 5 kHz and digitized at 10 kHz using a Digidata 1440A device (Molecular Devices). Data analysis was performed with Clampfit 10.2 (Molecular Devices) software.

### Plant materials and growth conditions

The Columbia-0 ecotype of *Arabidopsis thaliana* was the wild type used in this study, and the genetic backgrounds of the mutant and transgenic lines employed were as follows: *abcb4-2*, a transfer DNA insertion allele with a null phenotype^36^ and the *eir1-1* alele, a R1013K substitution in *PIN2* caused by a G-to-A mutation at position +3038 (Roman et al., 1995). The *eir1-1;abcb4-2* double mutant was generated by crossing *abcb4-2* into *eir1-1* and a line homozygous for both mutations was verified using PCR. WT primers for ABCB4 genomic DNA were as follows: 5’- GCGCAATACCTCTTTGGTTCATTAACT-3’ and 5’- GCGCATCATCCAACACTCTTCCTGATT-3’. The T-DNA Lb1a primer 5’- TGGTTCACGTAGTGGGCCATCG-3’ and ABCB4 genomic DNA primer 5’- GCGCAATACCTCTTTGGTTCATTAACT-3’ were used to screen for *abcb4-2*. Derived cleaved amplified polymorphic sequence (dCAPS) PCR was used to screen for *eir1-1*. Primers were as follows: 5’-TGATGTTGTTGATCATTTTATGGGACC-3’, which introduced an AgeI restriction site in WT EIR1 gDNA but not *eir1-1*, and 5’- CCTTAGGGCCATCGCAAACCC-3’. Resulting PCR products were digested with AgeI to identify lines that were homozygous for *eir1-1*. Seeds were sown on the surface of petri plates containing 0.8% phytoagar supplemented with one-half-strength Murashige and Skoog (MS) medium containing 2.15 g L^−1^ MS nutrient mix (Sigma-Aldrich), 1% (w/v) Sucrose, and 0.5 g L^−1^ MES, adjusted to pH 5.7 with KOH. For gravitropism studies, plates containing seeds were maintained at 4°C for at least 2 d. After this stratification treatment, plates were placed vertically at 23°C under a 16-h-light/8-h-dark photoperiod for 96 h.

### Gravitropism and growth rate

Seedlings for these assays were grown on petri plates containing the media described above. Plates were rotated 90° with respect to the gravitational vector and digital images were automatically collected every 2 min for 8 h using a bank of CCD cameras and infrared backlighting as described previously (Durham Brooks et al., 2010). The image files were automatically analyzed to calculate root tip angle time courses and growth rates with algorithms similar to those Durham Brooks et al. (2010) used.

### FRET

A Zeiss LSM 780 Meta confocal imaging system with a 30-mW argon laser and a 63X 1.4–numerical aperture oil immersion Plan-Apochromat objective was used to visualize live HEK293T cells or *N. benthamiana* epidermal cells coexpressing ABCB4 or PIN2 that were C-terminally tagged with CFP for ABCB4 or YFP for PIN2. FRET was measured by acceptor photobleaching (Herrick-Davis et al., 2006), with the following modifications. Prebleach CFP and YFP images were collected simultaneously following excitation at 458 nm (15% laser intensity). A selected region of interest was irradiated with the 514-nm laser line (100% intensity) using a 458-nm/514-nm dual dichroic mirror for 5 to 10 s to photobleach YFP. Postbleach CFP and YFP images were collected simultaneously immediately following photobleaching. Using ZEN vX software, FRET efficiency was measured as an increase in CFP fluorescence intensity from the ROI following YFP photobleaching and compared to an ROI selected from the background in order to account for noise. The FRET ratios at all the pixels within the region of interest were averaged to quantify the interactions of ABCB4 and PIN2 as done previously for glutamate receptor ion channel subunits (Vincill et al., 2013).

## Acknowledgements

This work was funded by NSF grant IOS-1360751 to E.P.S.

## Author Contributions

Edgar Spalding and Stephen Deslauriers designed the research; Stephen Deslauriers performed the experiments; Stephen Deslauriers and Edgar Spalding analyzed the data; Edgar Spalding and Stephen Deslauriers wrote the paper.

---

**Supplemental Figure 1.** Tryptophan and benzoic acid are not transported by ABCB4 or PIN2 according to reversal potential analysis. A, Average E_rev_ values obtained in each of the indicated conditions. B, The change in E_rev_ due to changing intracellular 0.1 μM IAA with each of the indicated test substrates. Only when 0.1 μM IAA was changed to 1 mM IAA was the change in E_rev_ statistically significant, as indicated by the asterisks (p=0.05).

**Supplemental Figure 2.** Confocal fluorescence micrographs of live HEK cells expressing ABCB4-CFP (a,e) and PIN2-YFP (b,f) before (a,b) and after (e,f) the region of interest marked by a red line was photobleached. Overlaying the CFP and YFP panels (c,g) shows an increase in CFP fluorescence following photobleaching, indicative of pre-bleaching quenching of ABCB4-CFP fluorescence by PIN2-YFP protein. Panels d and h show an enlarged view of the region of interest before and after photobleaching of the YFP.

**Supplemental Dataset 1.** All of the current-voltage (I-V) curves are contained in one comma separated value (csv) file. The results are grouped by the figure in which the mean values appear. All of the electrophysiology results in the paper can be reconstructed from these patch clamp trials.

**Supplemental Dataset 2.** All of the individual FRET efficiency values used to create the means in Fig. 5 are assembled in on comma separated value file. The file also contains the donor (CFP) and acceptor (YFP) fluorescence intensities obtained before and after photobleaching of the acceptor fluorophore.

